# Towards an Ecological Trait-data Standard

**DOI:** 10.1101/328302

**Authors:** Florian D. Schneider, Malte Jochum, Gaëtane Le Provost, Andreas Ostrowski, Caterina Penone, David Fichtmüller, Anton Güntsch, Martin M. Gossner, Birgitta König-Ries, Pete Manning, Nadja K. Simons

## Abstract

1. Trait-based approaches are widespread throughout ecological research, offering great potential for trait data to deliver general and mechanistic conclusions. Accordingly,a wealth of trait data is available for many organism groups, but, due to a lack of standardisation, these data come in heterogeneous formats.
2. We review current initiatives and infrastructures for standardising trait data and discuss the importance of standardisation for trait data hosted in distributed open-access repositories.
3. In order to facilitate the standardisation and harmonisation of distributed trait datasets, we propose a general and simple vocabulary as well as a simple data structure for storing and sharing ecological trait data.
4. Additionally, we provide an R-package that enables the transformation of any tabular dataset into the proposed format. This also allows trait datasets from heterogeneous sources to be harmonised and merged, thus facilitating data compilation for any particular research focus.
5. With these decentralised tools for trait-data harmonisation, we intend to facilitate the exchange and analysis of trait data within ecological research and enable global syntheses of traits across a wide range of taxa and ecosystems.

## Introduction

Functional traits are phenotypic (i.e. morphological, physiological, behavioral) characteristics that are related to the fitness and performance of an organism (McGill et al. 2006; Violle et al. 2007). Because trait-based approaches allow studying both patterns and mecanisms (Lavorel and Garnier 2002; Díaz et al. 2016), recent years have seen a proliferation of trait-based research in a wide range of fields. Trait-based studies have been conducted in a wide range of thematic areas ranging from the evolutionary basis of individual-level properties (Salguero-Gómez et al. 2016) to global patterns of biodiversity (Díaz et al. 2016) and ecosystem functioning (Bello et al. 2010; Allan et al. 2015). The trait framework relates losses of ecosystem function to changes in the functional composition of species assemblages (Mouillot et al. 2013; Perović et al. 2015). This offers the mechanistic background to relate biodiversity to climate change or local anthropogenic land use (Díaz et al. 2011; Lavorel and Grigulis 2012; Allan et al. 2015). Using traits is also a promising means of bypassing taxonomic impediment, i.e. the fact that a majority of species are yet undescribed and little is known of their interactions with the environment and other organisms. This is because functional traits allow us to infer the ecological role of organisms from their apparent features, regardless of their taxonomic identity (Duarte et al. 2011; Schrodt et al. 2015; Le Provost et al. 2017).

Many issues in trait-based research arise when compiling datasets from several sources. Data may differ in taxonomic nomenclature and resolution (e.g. reported on species level or aggregated on higher taxonomical orders), the scale and place of the study context, or the accurracy of the methodology applied in measurements. These differences are not always documented in the metadata accompanying a dataset. All of these factors render trait data extremely heterogeneous and make the task of data compilation time-consuming or even prohibitive. However, fully exploiting the potential of trait-based approaches relies heavily on the broad availability and compatibility of trait data to achieve sufficient taxonomic and regional coverage, both of present-day taxa ase well as in evolutionary deep-time.

To this end, the number of available trait datasets is increasing rapidly. In the past, trait data have been standardised and compiled in centralised databases for specific organism groups and regional scope, often centred around particular research questions (e.g. Pan THERIA, Jones et al. 2009; TRY, Kattge et al. 2011a; AmphiBio, Oliveira et al. 2017).These initiatives map heterogeneous data into a common scheme and, importantly, also offer access control and data usage policies. As such, they protect the rights of the original data providers while simplifying data queries for synthesis researchers. Besides initiatives aiming at assembling data, other tools to enable the compatibility of data across databases are being developed. These include semantic-web standards (Page 2008; Wieczorek et al. 2012) and ontologies of standard terms (Walls et al. 2012; Garnier et al. 2017). Meanwhile, open-science reaches the mainstream: it has become the declared goal of an open biodiversity knowledge management (http://www.bouchoutdeclaration.org/) and is increasingly demanded by journals and public research funding (German Science Organisaions 2010; Centre 2012; Swan 2012; Allison and Gurney 2015; Emerson et al. 2015). As a result, an increasing number of individual research projects publish their primary data on file hosting services like Figshare.com, Dryad (datadryad.org), Researchgate.net, or Zen odo.org, where no data standards are forced upon the uploaded material. It is likely that trait data will become increasingly available, but a lack of data and metadata standardisation will hamper the efficient re-use and synthesis of published datasets.

In this paper, we review existing trait databases and online portals, as well as initiatives for standardisation. We discuss current practice and the importance of data standards for trait-based research, and we identify current deficits in standardisation from a pragmatic view of data providers and data users. Based on these considerations, we propose a minimal structure and vocabulary for describing trait datasets, that builds upon and is compatible with existing terminology standards for biodiversity data. Finally, we present an R package that assists the harmonisation of trait data from distributed sources. With this easy-to-useterminology and toolset, we hope to convince trait-data providers and trait-data users aboutthe general importance of trait-data standardisation and lay out the roadmap towards an accessible ecological trait data standard.

### A review of initiatives for trait-data standardisation

In this section, we review four types of initiatives that are of relevance for trait-data standardisation (see Glossary in Table 1 for italicised terms):

**Table 1:** Glossary of terms from the biodiversity data-management context as they are used in this paper; draws from Garnier et al. (2017).

| Term | Definition |
| --- | --- |
| Term | A word that describes a particular *concept* as part of the specialised vocabulary of a field |
| Concept | An idea, notion or object that is made explicit in an information context by *name*, definition, *URI* or other reference (https://www.w3.org/TR/skos-reference/) |
| Controlled vocabulary | A list of *terms* that gives all valid consensus terms for a particular context, while no unlisted entries are accepted |
| Terminology | The body of *terms* and *concepts* used with a particular application in a subject of study, usually formalised in a *thesaurus* or *ontology* |
| Data standard | A published set of instructions and *terminologies* for storing and exchanging data content of a particular type (e.g. trait data), that is recognised by a large proportion of members of the application context |
| Thesaurus | *Controlled vocabulary* that provides key *terms* with their associated *concepts* for a specific field or domain of interest (Laporte et al. 2013) |
| Ontology | *Controlled vocabulary* that (opposed to a *thesaurus*) relates *concepts* to each other by cross-references, e.g. defines a hierarchy of terms; thus a formal model of the objects and their relationships in a domain of interest (Gruber 1995) |
| Semantic web | An extension of the world wide web that aims for machine-readable meaning of information via well-defined *data standards*, *ontologies* and exchange protocols (Berners-Lee et al. 2001); the World Wide Web Consortium (W3C) defines standards, i.e. specifications of protocols and technologies for the semantic web (http://www.w3.org/standards/semanticweb/) |
| Dataset | A set of measurements and observations; often originating from a single experimental set-up or study context; can be considered as being internally homogeneous across all data entries |
| Data table | A two-dimensional spread-sheet containing data organised in rows and columns; in most cases these data are considered 'static', i.e. they are not altered or filtered across time |
| Database | A suite of data compiled from multiple *datasets*, i.e. from multiple study contexts or observation types; may take the form of a two-dimensional data table, but mostly is organised in into *relational databases* using database software; |
| Relational database | Usage in this paper: Two or more *data tables* that are related by common information contained in one or more columns; common information is usually labelled by *identifiers* (IDs) |
| Online database | A *relational database* that is made accessible on the internet; offering forms for filtering and downloading subsets of the data; some online databases offer access via a *webservice* and an *API* that can be addressed computationally |
| Identifier (ID) | A unique label that relates entries within and across *datasets*; is used to connect *data tables* into a *relational database*; can be user-specific or, as a *URI*, point to a globally valid *ontology* or *thesaurus* |
| Uniform Resource Identifier (URI) | An unambiguous pointer to a unique resource on the internet; used to refer to single terms of a *thesaurus* or *ontology*; an example is 'http://t-sita.cesab.org/BETSI_vizInfo.jsp?trait=Body_length' |
| Webservice | An exchange protocol to access *online databases* directly and programmatically, i.e. by calls from a software tool |
| Application Programming Interface (API) | A set of clearly defined methods of communication between software components, e.g. client software and the *webservice* of an online databases; APIs are usually documented on the website of a database provider |
| Online Portal | A website designed as a platform for the exchange of information, e.g. trait data; a portal may include a communication forum, data upload forms, a database access point, and advanced user management for data access |
| File repository | A short-term storage of *datasets* or long-term archiving on *file-hosting services*; online repositories make data available for public access, provide *metadata* and (not always) facilitate citations via DOIs (Digital Object Identifiers) |
| File-hosting service | An online platform that hosts *datasets* or entire *repositories* and provides access to a wide audience on the internet; examples in biology are Figshare.com, Dryad (datadryad.org), Researchgate.net, or Zenodo.org |
| Metadata | Data documentation of the higher level information or instructions; describe the content, context, quality, structure, provenance and accessibility of a data object (Michener et al. 1997) |
| Darwin Core Standard (Dwc) | Body of *terminologies* providing terms intended to facilitate the sharing of information about biological diversity (http://rs.tdwg.org/dwc/) |
| Darwin Core Archive (DwC-A) | A file archive, or *repository* structure, that contains *metadata* (specified using Ecological Metadata Language, EML) and primary data combined into a *relational database* via *identifier* columns. |
| Method handbook | A listing of consensus methodology that is to be applied to acquire a particular measure, thus formalising the precise *concepts* of measures. |
| Occurrence | A single observation instance of a taxon, i.e. an organism at a particular place at a particular time (http://rs.tdwg.org/dwc/terms/Organism) |

1. Initiatives that provide *trait datasets* which have been assembled out of a particular research interest, either by measurement or collated from the literature.
2. Initiatives that aim to harmonise trait data from the literature or from direct measurement into *trait databases* and make those data widely available.
3. Initiatives that aim at the standardisation and development of consensus measurement methods and definitions for traits, and provides standard *terminologies* in the form of *thesauri* and *ontologies*.
4. Initiatives that aim to leverage *relational database* structures and *semantic web* technology to link trait data to a wider set of biodiversity data.

We discuss these initiatives separately although often they are developed in conjunction to serve a particular database project, as for instance in the case of the TRY plant database (Kattge et al. 2011a; Kattge et al. 2011b) and the Thesaurus of Plant Traits (TOP; Garnier etal. 2017). We show how the degree of trait-data standardisation in existing datasets spans this entire spectrum and which tools and standards are applied to achieve harmonisation of data from multiple, distributed sources. The objective of this review is to raise awareness for the generic structure of trait data and aid researchers to share and publish own datasets in an appropriate form.

### Trait datasets

In the field of comparative biology, morphological traits related to plant flower, leaf and stem traits or bird wing and beak measurements, as well as life-history traits such as Ellen-berg values for plants or ecological parameters of animals (e.g. reproductive traits, feeding biology, dispersal or body size) have been measured for decades, and have been published in regular journal articles or books. With the rise of ecological trait-based research, individual measurements and information available from species descriptions have been compiled into project-specific datasets that typically comprise a local set of taxa and a focal set of traits. A plethora of such static datasets has been published along with scientific articles or as standalone data publications (see Kleyer et al. 2008 for a review on plant data; on animal data, see e.g., Gossner et al. 2015; Ricklefs 2017). Today, the online publication of such data is greatly facilitated by *file hosting services* (e.g. Figshare.com), which warrant long-term accessibility and citability via DOIs, and support Public Domain dedication or Creative Commons licenses. These platforms offer publicly accessible repositories at low-cost or for free, which makes them attractive for small and intermediate sized research projects that cannot dedicate extra resources for data management. However, although open for manual access, the trait datasets on data repositories might be stored in proprietary (e.g. .xlsx, .docx) or binary (e.g. .pdf) data formats which make a programmatical extraction tedious and dependent on commercial software, putting the long-term and open accessibility of these data at risk. Most importantly, these platforms enable public hosting of data with very low thresholds for *metadata* documentation and data standardisation.

For trait data, there are typical issues arising from the variability of data structures. For instance, the column descriptions and terminology applied to taxa and traits are mostly project specific, and rarely chosen to allow translation into larger database initiatives. Fur-thermore, metadata varies in its detail, e.g. for documenting descriptions of variables, measurement procedures or sampling context (Kattge et al. 2011b). In terms of structure, trait data usually are reported in a species*×*traits wide-table format. In this format, each row contains a species (or taxon) for which multiple traits are reported in columns. Similarly, when reporting raw data, researchers place observations of individual organisms in rows with multiple trait measurements applied to the same individual across multiple columns. Variability in the number and meaning of columns in these *data tables* requires tedious manual adjustments when merging multiple datasets (Wickham 2014).

A global overview of existing trait data for all taxa and trait types is difficult to obtain. Therefore, in an attempt to collate a list of existing distributed datasets, we initiated a living spreadsheet (https://goo.gl/QxzfHy) which lists published trait datasets, their regional and taxonomic focus, the number and scope of traits covered, their location on the internet and the terms of use (see Appendix A for a current excerpt of this list). We invite data owners and users to add further trait datasets to this spreadsheet.

As it stands, the decentralisation and the lack of data standardisation of low-threshold online repositories renders the compilation of data into larger collections inefficient and reduces the potential of many published datasets to be re-used and combined into broad synthesis analysis.

### Database initiatives

In the past two decades, many distributed trait datasets have been aggregated and haronised into greater collections with particular taxonomic or regional focus (e.g. Klotz et al. 2002; Kleyer et al. 2008; Jones et al. 2009; Kissling et al. 2014; Myhrvold et al. 2015; Iversen et al. 2017; Oliveira et al. 2017, see Appendix A table A1). While mostly concerned with issues of heterogeneity in units or factor levels, and aiming for high taxonomic coverage, few of these datasets apply a standardised terminology for taxa or traits that would allow them to be efficiently related to other databases. Documentation of metadata and methodology differs in the level of detail, depending on the research focus of the initiative. Just as the individual datasets described above, many of these databases are published as static data tables on low-threshold file hosting platforms and are updated irregularly.

As they deal with much larger amounts of data, initiatives that form around natural history museum collections are more concerned with standardisation. Concerning organism traits, with the digitisation efforts that are currently undertaken in many museum collections (Vollmar et al. 2010; Blagoderov et al. 2012), supported by citizen science crowdsourcing (e.g. www.markmybird.org), data on body measurements are likely to grow exponentially in the near future. For example, the VertNet database compiled and harmonized large quantities of vertebrate trait data with the aim of mobilising measurements from collections (Guralnick et al. 2016). The resulting data are published as versioned data tables which are updated as new data sources become available.

More specialised trait-database platforms have been created to cover certain trait types (e.g. floral traits, seed traits, root traits or wood density traits), interaction types (e.g. pollination traits or feeding relationships), or a specific environmental and experimental context of the trait observation (e.g. location or climatic data). Such database initiatives attract data submissions from a defined research field and take care of the harmonisation process and thereby greatly facilitate data synthesis. For example, by aiming for a universal framework for plant traits, the TRY database (Kattge et al. 2011a) attracted more data submissions and downloads than any other trait data platform. The *online database* enables selective data download and user permission and rights management. As a community effort, TRY serves as a network for consensus building on trait definitions (Garnier et al. 2017) and measurement methodology (Perez-Harguindeguy et al. 2013) (see next section). Microbial ecologists also make frequent use of trait-based approaches to assess genomic function and describe functional diversity at the community level (Fierer et al. 2012; Fierer et al. 2014; Krause et al. 2014). Here, ‘operational taxonomic units’ (OTUs)are derived from metagenomic analysis (Torsvik and Øvreås 2002; Langille et al. 2013). Databases are also used to interpret OTUs in terms of their functional role (e.g. the KEGG orthology, Kanehisa et al. 2012). For animals, a single unified platform and harmonizing scheme for animal trait data is still lacking. The reason for this may be that harmonizing trait data on animals, which span multiple trophic levels and possess diverse body plans, is a more complex task than for plants (Moretti et al. 2017). Nonetheless, initiatives for particular groups of animals, such as the BETSI database collects traits on soil invertebrates (http://betsi.cesab.org/; Pey et al. 2014), and the Carabids.org web portal collects traits of carabid beetles (http://www.carabids.org/), already exist.

Regarding open access, few of these centralised databases comply with the criteria demanded by journals and funding agencies for primary data publication. The platforms incentivise data submissions by offering increased data visibility and usage, while providing data use policies that secure author attribution and potentially co-authorship. With the proactive turn towards open access data (as stated in the Bouchot Declaration; http://www.bouchoutdeclaration.org/), it may be necessary to find other incentives for data submission.

### Thesauri and Ontologies for traits

A major challenge in trait-data standardisation is the lack of widely accepted and unambiguous trait definitions. Previous standard definitions of trait *concepts* range from listings of selected definitions in *glossaries*, over well-defined methodological *handbooks* and comprehensive *thesauri*, to relational definitions of trait concepts in *ontologies*. While glossaries may be seen as specific for a study context, the initiatives behind method handbooks, the sauri and ontologies are primarily concerned with consensus building on trait definitions in a wider community.

Very general classes of traits are defined within the list of GeoBON Essential Biodiver sity Variables (Pereira et al. 2013). Assigning a more detailed and unambiguous methodological protocol to a trait, including the units to use or the ordinal or factor levels to be assigned, is key for standardising the physical process of measuring. Efforts to develop handbooks for measurement protocols provide such a methodological standardisation for plants (Cornelissen et al. 2003; Perez-Harguindeguy et al. 2013) or invertebrates (Moretti et al. 2017), but obviously are of limited use in harmonising trait data that pre-date or ignore this standard (Kattge et al. 2011b).

A thesaurus provides a “controlled vocabulary designed to clarify the definition and structuring of key terms and associated concepts in a specific discipline” (Laporte et al. 2013; Garnier et al. 2017). Expanding on this, ontologies link the defined terms by formally defining the relationships between them, with the objective of enabling a computational interpretation of data. Being publicly available, it is also possible to refer to these defined terms via globally unique *Uniform Resource Identifiers (URIs)* within own datasets. For example, a measurement of seed size could be linked to the Planteome Trait Ontology (TO) definition of ‘seed size’ by referencing ‘http://browser.planteome.org/amigo/term/ TO:0000391’. Ontologies define terms based on other well-defined terms from published ontologies. The TO definition of the concept ‘seed size’ contains references to other globally defined terms: “A seed morphology trait (TO:0000184) which is the size of a seed (PO:0009010).” Furthermore, trait definitions may refer to related terms or synonyms defined in other trait ontologies or other scientific ontologies, like units as defined by the Units of Measurement Ontology (Gkoutos et al. 2012). This way, each trait definition may link to a broader or narrower term. For example, the definition of ‘femur length of first leg, left side’ is narrower than ‘femur length’ which is narrower than ‘leg trait’ which is narrower than ‘locomotion trait’. By providing this interlinkage of trait ontologies, a machine-readable web of definitions is spun across the Internet which allows researchers and search engines to relate independent trait measurements with each other and connect it to the wider *semantic web* of online data (Berners-Lee et al. 2001; Page 2008). The distinction of thesauri and ontologies is not truly binary. Rather they mark idealised ends of a spectrum. While thesauri may contain defined relations between terms within the standard, ontologies relate most terms to other defined concepts, and also link those to other standards.

Comprehensive trait thesauri have been developed in the TOP Thesaurus of plant traits, which is employed in the TRY database (Garnier et al. 2017), and in the Thesaurus for Soil Invertebrate Trait-based Approaches (T-SITA, http://t-sita.cesab.org/, Pey et al. 2014). Ontologies of trait definitions have been developed for plants (e.g. the Plant Ontology, Jaiswal et al. 2005; the Flora Phenotype Ontology, Hoehndorf et al. 2016), as well as for animals (e.g. the Hymenoptera Anatomy Ontology, Yoder et al. 2010; the vertebrate trait ontology, Park et al. 2013). The existing thesauri and ontologies for traits differ widely in terms of hierarchical depth and detail, as well as in curation efforts and measures for peer-reviewed quality control. Meta-ontology initiatives, like Planteome.org, offer access to multiple published ontologies and build platforms for their collaborative development (Walls et al. 2012). For general biodiversity data, the OBO Foundry (http://www.obofoundry.org/), Ontobee (http://www.ontobee.org/), Bioportal (https://bioportal.bioontology.org/), or the GFBio Terminology service (https://terminologies.gfbio.org/), provide centralized hosting for advanced trait ontologies and offer webservices for computational access.

To conclude, there is already a suite of globally available thesauri and ontologies for traits that emerged from standardisation efforts of methodologies and community consensus processes. However, definitions in some domains are better covered than others. Interlinkage and accessibility of ontologies can be much improved to fulfil semantic web standards. Most importantly, while these defined vocabularies are widely used in biodiversity data management, distributed data repositories of smaller project contexts hardly make use of them. A more widespread implementation of ontologies would advance the possibilities to aggregate datasets into databases and reduce noise and uncertainty. To achieve this, the use of ontologies and thesauri must be incentivised and facilitated for individual researchers. For example, the accessibility of ontologies will increase if open *Application Programming Interfaces (APIs)* are provided as a way to extract the definitions and higher-level trait hierarchies programmatically via software tools. Software then can assist researchers in linking own data to globally defined concepts.

### Trait-data structures for the semantic web

While trait thesauri and ontologies typically define traits for focal groups of organisms, they do not specify the format or structure in which trait data should be stored and linked to further standard terminologies, such as standard taxonomy nomenclatures.

To make sense of trait data in the context of more general databases, a consensus definition of trait data is necessary.

Trait data have been defined by Garnier et al. (2017) to follow an entity-quality model (EQ), where a trait observation is ‘an entity having a quality’. More specifically, a trait dataset contains information on quantitative *measurements* or qualitative *facts* (i.e. trait values) describing the physical phenotypic characteristics relating to fitness and performance (i.e. traits) observed on a biological entity (i.e. an individual specimen, or parts of an individual specimen) that can be assigned to a biological taxon (i.e. a species or higher-level taxon). We are expanding on this definition: quantitative measurements are values obtained either by direct morphological, physiological or behavioural observations on single specimens (Fig. 1*a*), by aggregating replicated measurements on multiple entities (Fig.1*b*) or by estimating the means or ranges for the respective taxon as reported in the literature or other published sources (e.g. databases, Fig. 1*c*). Qualitative facts are assignments of an entity to a categorical level, e.g. of a behavioural or life-history trait (Fig. 1*d*). The entity or observation (i.e. the *occurrence*) to which the reported measurement or fact applies may differ in organisational scale – depending on the scientific question – and could be a sub-sample or bodypart, an individual specimen, an entire species or a higher-level taxon (e.g. a genus).

**Figure 1:**
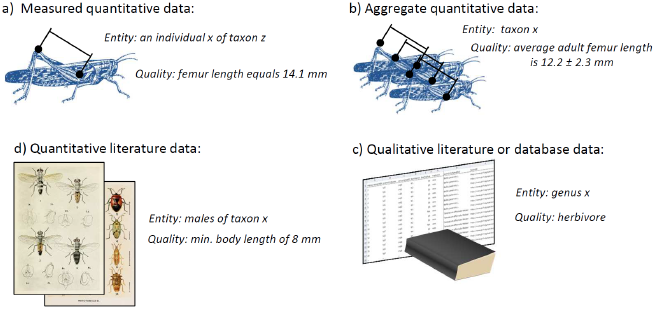
Types of ecological trait data assume different entities or reported quantities. a) morphometric or morphological measurements of individual body features (lengths, areas, volumes, weights) or other quantities related to life history (e.g. reproductive rates, life spans); b) aggregated traits are reported as means taken on multiple measures of members of a taxon; c) quantities may be extracted from literature or existing databases, referring to the entire taxon (or a subset, e.g. a sex) as the entity of description; d) Qualitative traits are categorical or binary descriptors of the entire species or higher taxonomic group.

These relationships between a trait observation and an individual organism as an occurrence of a particular taxon have been formalised in the schema for biological collection records (ABCD Schema; Holetschek et al. 2012) and the Darwin Core Standard for biodiversity data (DwC; Wieczorek et al. 2012). For example, the Global Biodiversity Information Facility (GBIF, www.gbif.org) applies these terms. These frameworks specify terms and classes to describe the general structure of biodiversity databases, for example by defining names for columns that contains measurement values, units, taxon names, variables such as sex or life stage, ancillary information of time and date of observation, and methodological details. The terminologies provided by these standards are quite universal and even cover most use cases of trait data. An entire ecosystem of data standards links to and expands the capacities of DwC (Wieczorek et al. 2012).

Specifically designed for plant traits, Kattge et al. (2011a) proposed a generic database structure that covers most potential use cases of trait-based ecology. This data structure is built around a central data table that contains observations, i.e. a single event of measurement on the same individual plant specimen at the same point in time. This structure emphasises the fact that multiple trait data are measured on the same individual organisms and used to analyse correlations between these multiple traits. Identifiers link the measurements (qualities) to the same observation (entity), each measurement being well defined by additional standard tables. The observations are also linked to a taxonomy and ancillary descriptors of the observation context, like location or experimental treatment. This structure can be implemented in any relational database management system.

In a similar vein, the Encyclopedia of Life (EOL) project has proposed the database framework TraitBank (Parr et al. 2016) for major physiological and life-history traits of all kingdoms of life, which is to date the most general approach of an integrated structure for trait data. The framework employs established terms provided by the DwC, relates trait definitions to trait ontologies for phenotypic or anatomical terms, and maps taxa to global identifiers in taxonomic hierarchies of name service providers to capture synonyms, misspellings and controversies (Parr et al. 2016, http://eol.org/info/cp_archives). Additional layers of information capture bibliographic reference, multimedia archives and ecological interactions. TraitBank invites data submissions to the EOL database in a structured Darwin Core Archive (DwC-A, Robertson et al. 2009), a zip-file with annotated text-files that is also preferred for observation data in GBIF (GBIF 2017, http://tools.gbif.org/dwca-assistant/). The archive also integrates the general framework for metadata of the Ecological Metadata Language (EML, KNB 2011). The difficulties with keeping taxonomic references intact along with continuous changes in taxonomy consensus are a central challenge of biodiversity data management and are beyond the scope of this review (Franz et al. 2016). Initiatives that aim at providing a stable reference for taxa are for instance the EOL Catalogue of Life (http://www.catalogueoflife.org/, Roskov et al. 2018), the GBIF Backbone Taxonomy (Secretariat 2017), or the EDIT Platform for Cybertaxonomy (https://cybertaxonomy.eu/).

These proposed standards are responses to a demand from biodiversity data managers for more structured input from the research community. However, hardly any of the aforementioned trait datasets for birds, amphibians, or mammals employs such ontologies or semantic web standards. One reason for this is most certainly complexity: the data structures are designed for multi-layered, relational databases rather than for standalone datasets for which a two-dimensional data table may suffice. In the eyes of the data-provider, in most cases, ancillary co-factors can be appended as extra columns to the dataset. The other reason is lack of awareness for the need for trait-data standardisation among data providers: many providers are not trained in the demands of biodiversity data-management and complying with what may be non-intuitive data structures is an investment without clear incentive or immediate pay-off, and hardly affordable for small and intermediate-size research projects.

By filling this gap, data-brokering services (e.g. the German Federation for Biological Data; gfbio.org; Diepenbroek et al. 2014) or data management systems for scientific projects (e.g. KNB and its open-source database back-end Metacat, https://knb.ecoinformatics.org/; Diversity Workbench, www.diversityworkbench.net; BExIS, http://bexis2.uni-jena.de/;) are likely to gain importance. These services simplify and direct the standardised upload of research data and descriptive metadata into reliable and interlinked data infrastructures. One goal of such initiatives is to facilitate data publications and standardisation for researchers, for instance by providing terminologies and ontologies for biodiversity data, and by consulting on publication licenses.

### Conclusion of review

Initiatives for standardisation (e.g. ontologies and data standards) and platforms for data management (e.g. database and data management platforms) provide great visibility and improve interconnectedness of datasets, but raise relatively high thresholds for data and metadata preparation. Low-threshold repositories offer the hosting of scientific primary data attracting a wealth of heterogeneous trait datasets, but data harmonisation of these distributed data sets is currently laborious. The goal must be to better integrate these distributed data into the global biodiversity data-management ecosystem by creating awareness for data standardisation on the side of data providers. We propose the development of tools and vocabularies that impose low thresholds and offer high pay-off in the visibility and interconnectedness of published data.

## An ecological trait-data standard vocabulary

As a response to the challenges outlined above, we propose a versatile vocabulary for trait-based ecological research. The aim of the vocabulary is to cover the variety of trait-based approaches and their different degrees of measurement detail. Rather than describing a data structure for relational databases, the vocabulary is intended as a more inclusive terminology, that can be used in simple two-dimensional datasets as well as in the exchange of data between web services in the semantic web. By using this standard vocabulary, authors can ensure that the description of trait measurements that are uploaded to distributed data repositories will be unambiguous and generally applicable. It will facilitate re-use of data for future data aggregation initiatives and data synthesis and ensure long-term accessibility.

In designing this vocabulary, we drew on the combined expertise of empirical biodiversity researchers (data providers), biodiversity synthesis researchers (data users), and biodiversity informatics researchers (data managers). We paid particular consideration to the work of Kattge et al. (2011a), Kattge et al. (2011b), and Garnier et al. (2017), as well as Parr et al. (2016) to ensure compatibility of our proposed data structure with major trait databases and existing standards for biodiversity data management. Here, the use of identifiers (‘IDs’) for the individual measurement observations (‘measurementID’),specimens(‘occurrenceID’), sampling events (‘eventID’), or taxa (‘taxonID’) is key to map two-dimensional data onto the structure of relational databases. Besides being used for the publication of datasets, the standard vocabulary could be imposed in webservices or download tools, e.g. APIs that provide direct access to online databases. The vocabulary proposed is intended to form the foundations of a standard nomenclature that can be expanded and corrected by the wider community of researchers using trait-based approaches in ecology.

### How to apply the standard vocabulary

We suggest that any trait dataset that is published on online repositories should draw its column names and field entries from the defined vocabulary where possible. The core vocabulary lists and defines terms that describe a dataset according to the Entity-Quality model described above (Garnier et al. 2017): each entry describes a trait value (i.e. quality) observed on an individual or population (i.e. entity), of a biological taxon. When applying the vocabulary, it is implicit to use a two-dimensional observation long-table format for the data (Fig. 2 *b*), rather than a species × traits matrix (Fig. 2 *a*). As the long-table format draws from a defined set of columns, merging datasets is easier. Long-table datasets also purport multiple advantages for data manipulation (e.g. filtering, sub-setting and aggregating data, Wickham 2014).

**Figure 2:**
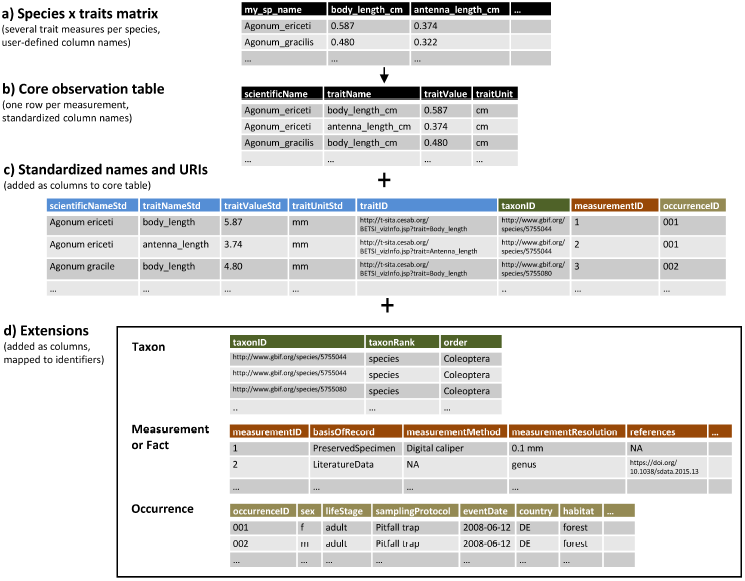
Formats used for trait datasets. a) taxon-level trait data compiled from literature or aggregated from measurements are often published as a compiled species × traits matrix; b) observation long-tables are a well defined and tidy data format, reporting one single measurement per row and c) relating it to a standard trait definition and accepted taxon name using unambiguous identifiers. Additional identifiers relate each row to other layers of information on d) the taxon resolution, the specimen (occurrence) or the origin or confidence on the reported measurement or fact.

Well-defined identifiers (‘IDs’) are key elements to structure the datasets and relate them to complementing datasets, if necessary (Fig. 2 *c* & *d*). For instance, for occurrence level data where multiple trait measurements are reported for each individual specimen, the same user-defined entry for ‘occurrenceID’ would link several measurements across the rows of the dataset. Similarly, multivariate measurements, for instance gas chromatography data or x-y-z data of morphometric landmarks could be linked via a ‘measurementID’. In literature data, summarised traits are usually given at the taxon level instead of the individual organism (e.g. reported as means or factorials) and a ‘taxonID’ is the key identifier. In larger compilations, a ‘datasetID’ allow to trace data origin to the primary source. Beyond being just of structural use for the dataset, identifiers are capable of linking own data to consensus taxonomy and trait terminology via URIs, which point to external terminology services (see above for resources). Two-dimensional spreadsheets are however limited in the number and complexity of co-variates they can contain. As such, for datasets containing multi-layered information on observations, traits, taxa and environmental context, the use of relational datatabase structures may be indicated, like the generic trait database structure proposed by Kattge et al. (2011b) or the TraitBank structure proposed by Parr et al. (2016). The trade-off is user-side readability and handling in a single table *vs.* avoidance of content duplication and redundancy in a relational database. The standard vocabulary proposed here may still be applied to describe columns within the individual data tables of relational databases.

For reasons of long-term accessibility, data should not be uploaded in proprietary spreadsheet formats (like ‘.xlsx’) but rather in comma-separated text files (‘.csv’ or ‘.txt’) that are compatible with all computing platforms and internationalisation settings by applying a unified character encoding (e.g. UTF-8 or ASCII).

In order to ensure traceability, the metadata of any dataset that employs this vocabulary should refer to the specific online version that was used to build the dataset, e.g.“Schneider, F.D., Jochum, M., Le Provost, G., Penone, C., Ostrowski, A. and Simons, N.K., 2018 Ecological Traitdata Standard v0.8, DOI: 10.5281/zenodo.1255287, URL: https://ecologicaltraitdata.github.io/ETS/v0.8/”. In addition to this versioned online reference, the dataset should also cite this paper for an explanation of the rationale. Wherever referring to individual terms of the vocabulary in publications or metadata, this should be done via their global identifiers, which will be hosted by the GFBio Terminology Service (Karam et al. 2016, https://terminologies.gfbio.org/) and can be accessed programmatically (i.e. via the API; **in preparation!**). Wherever our glossary refines or dupllicates existing terms from other ontologies for biological data, like the Glossary of EOL (http://eol.org/info/516) and Darwin Core (http://rs.tdwg.org/dwc/terms/), we indicate this in the fields ‘refines’ or ‘identical’, respectively.

### Terms of the standard vocabulary

The standard vocabulary is accessible at https://ecologicaltraitdata.github.io/ETS/. The core terms describe minimal trait data according to the Entity-Quality model. Beyond these core observations, further information might be available that are related to the taxonomic assignment, or that put the reported fact, measurement or sampling event in a broader observation context (including geolocation and date information). These information can be useful for future analysis of the causal reasons of trait variation and should always be published along with the core data. For this case, we offer three extensions of the core vocabulary (“Taxon”, “Measurement or Fact”, and “Occurrence”) that expand and refine terms of the Darwin Core Extensions (see below) which may simply be added as extra columns to the core dataset. Additional terms are provided for metadata and for relating trait names to definitions and external ontologies or thesauri (see section on metadata below). The scope of the vocabulary may not yet cover all aspects of morphological and evolutionary perspectives. Also, information about interactions between species are not within the scope of the Entity-Quality Model, but may easily be combined with trait data by using other extensions of DwC. Therefore, we invite researchers to contribute to the next iterations of the standard vocabulary and develop own applications and ontologies that interact with it.

### Specification of core terms

To qualify as trait data according to the definition provided above, where each row is the reported measurement or fact for a single observation, the following columns are required at minimum (Fig. 2 *b*): 1. a value (column `traitValue`) and – for numeric values – a standard unit (`traitUnit`); 2. a descriptive trait name (`traitName`) that links to a well-defined definition; 3. the scientific taxon name for which the measurement or fact was obtained (`scientificName`). For these core values, unambiguous and self-explanatory vocabularies for trait names and taxa are recommended. However, to ensure compatibility with existing databases or analytical code, it might be necessary to use abbreviations or user-specific identifiers for `scientificName` and `traitName` instead. In this case, it is essential to relate the user-defined names to a consensus standard of taxon names as well as a look-up table of traits. This is achieved by adding globally valid Uniform Resource Identifiers (URIs) for taxon (`taxonID`) and trait definitions (`traitID`), complemented by the human-readable verbatim accepted names (`ScientificNameStd` and `traitNameStd`, respectively). For example, referring to GBIF Backbone Terminology, for *Bellis perennis*, the taxonID would be ‘https://www.gbif.org/species/3117424’; the `traitID` for ‘fruit mass’ according to TOP Thesaurus of plant traits would be ‘http://top-thesaurus.org/annotationInfo?viz=1&&trait=Fruit_mass’.

By allowing for a double record of both user-specific and standardised entries, we acknowledge the fact that most authors have their own schemes for standardisation which may refer to different scientific community standards (as practised in TRY; Kattge et al. 2011a). This redundancy of data allows for continuity for data owners while also ensuring quality checks and comparability for the data user.

### Extensions for additional data layers

Beyond measurement units or higher taxon information, further information might be available that may not be core data, but are related to the individual or specimen, or to the reported fact, measurement or sampling event. The data standard provides three extensions of the vocabulary that should be used to describe this information (Fig. 2*d*):

- The Taxon extension provides further terms for specifying the taxonomic resolution of the observation and to ensure the correct reference in case of synonyms and homonyms. (http://ecologicaltraitdata.github.io/ETS/#extension-taxon)
- The `MeasurementOrFact` extension provides terms to describe information at the level of single measurements or reported facts, such as the original literature from where the value is cited, the method of measurement or statistical method of aggregation. It provides important information that allows for the tracking of potential sources of noise or bias in measured data (e.g. variation in measurement method) or aggregated values (e.g. statistical method applied), as well as the source of reported facts (e.g. literature source or expert reference). (https://ecologicaltraitdata.github.io/ETS/#extension-measurement-or-fact)
- The `Occurrence` extension contains vocabulary to describe information on the level of individual specimens, such as sex, life stage or age. This also includes the method of sampling and preservation, as well as date and geographical location, which provides an important resource to analyse trait variation due to differences in space and time. (https://ecologicaltraitdata.github.io/ETS/#extension-occurrence)

Many terms of these extensions refine or copy terms of the DwC and their own Taxon, MeasurementOrFact and Occurrence extensions and EOL TraitBank’s use of those terms (http://eol.org/info/structured_data_archives). These additional layers of information can either be added as extra columns to the core dataset or kept in separate data sheets (published separately or as part of a Darwin Core Archive), thus avoiding redundancy and duplication of content. A unique identifier would link to these other datasheets, encoding each individual occurrence of a species (`occurrenceID`), single measurements or reported facts (`measurementID`), locations of sampling (`locationID`) and sampling campaigns (`eventID`). Some data-types may directly refer to existing global identifiers for occurrence IDs, e.g. a GBIF URI or a museum collection code references the precise specimen from which the measurement was taken (Groom et al. 2017; Güntsch et al. 2017).

### Specification of Metadata

Wherever possible, the column `traitID` should point to a publicly available, unambiguous trait definition in a published ontology. If no globally available trait definition exists as an external reference, trait datasets should always be accompanied by a dataset specific list of traits as part of the metadata or as an accompanying data table. Such a controlled vocabulary would, in its simplest form, assign trait names with an unambiguous definition of the trait and an expected format of measured values or reported facts (e.g. units or legit factor levels). from published trait ontologies. By providing a minimal vocabulary for trait lists (see https://ecologicaltraitdata.github.io/ETS/#terms-for-trait-definitions), we hope to facilitate the unambiguous definition of traits for trait datasets. This vocabulary might also prove useful for the future publication of trait ontologies.

Information about the authorship and ownership of the data and the terms of use should be considered when sharing and working with trait datasets. We define a vocabulary (https://ecologicaltraitdata.github.io/ETS/#metadata-vocabulary) that allows trait data to be related to authors and owners, while also stating a bibliographic reference and license model. In the case of primary measurement data, this information applies to the entire trait dataset, and should be stored along with the published data as metadata (e.g. in a separate metadata file, possibly applying the ecological metadata language, EML). In cases where individual data from different sources are compiled into a trait database, these information must be provided at the measurement level. This can be achieved by appending the information as columns to the core dataset, or via an unambiguous `datasetID` and a descriptive `datasetName`.

## Computational tools for producing compliant data

To access data from public databases, the R-package ‘traits’ (Chamberlain et al. 2017) contains functions to extract trait data via several open API interfaces including Birdlife, EOL TraitBank or BetyDB. The package ‘TR8’ provides similar access to plant traits from a list of databases (including LEDA, BiolFlor and Ellenberg values; Bocci 2015) and aggregates them into a species × traits matrix. However, none of these packages provide the option to harmonisation trait data into a unified scheme. To close this gap, we developed the R package ‘traitdataform’, which assists the production of data compliant with the trait data standard proposed above. There are two major use cases for the package:

1. preparing trait datasets for publication on public hosting services and project databases, and
2. automating the harmonisation of trait datasets from different sources by moulding them into a unified format.

A comprehensive documentation of the package can be found on its Github repository (https://github.com/EcologicalTraitData/traitdataform) and the documentation website (http://EcologicalTraitData.github.io/traitdataform/). The package is under continuous open source development and invites participation in development, comments or bug reports via the Github Issue page (https://github.com/EcologicalTraitData/traitdataform/issues).

The key function of the package is `as.traitdata()` which moulds a species-traitmatrix or occurrence table into a measurement long-table format (Fig. 3). This function also maps column names into terms provided in the trait data standard and adds metadata as attributes to the output object. This example converts an own file ‘data.csv’ into a dataset of long-table structure that employs the standard vocabulary for core data:

**Figure 3:**
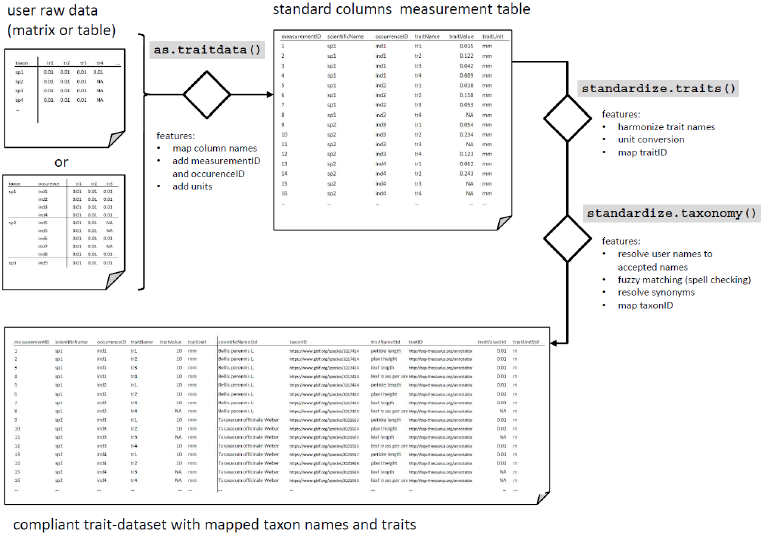
Process chart of the functions provided within the R package ‘traitdataform’ to apply the standard vocabulary to any trait-data table.

~~~
library(“traitdataform”)
dataset <- as.traitdata(read.csv(“path/to/data.csv”),
  traits = c(“body_length”, “antenna_length”,
                     “metafemur_length”),
  units = “mm”,
  taxa = “name_correct”,
  keep = c(locationID = “location”)
  )
~~~

The parameter ‘traits’ lists column names that contain trait values. The column containing taxon names is given in parameter ‘taxa’. Note that the parameter ‘keep’ specifies and renames any data that should be maintained in the output. The parameter ‘units’ is used to specify the input units of measurement. In order to map user-provided names to unambiguous and globally unique identifiers, the function `standardize.taxonomy( )` matches scientific taxon names automatically to the GBIF Backbone Taxonomy and adds the column `taxonID` to the core data (Fig. 3).

The R-package further supports the mapping of trait names to a list of trait definitions and identifiers (this lookup table is cast into an own object class called ‘thesaurus’). The following example harmonises traits based on a minimal list, referencing trait names with globally valid URIs provided by the BETSI thesaurus of soil invertebrate traits:

~~~
traitlist <- as.thesaurus(
 body_length = as.trait(“body_length”,
 expectedUnit = “mm”, valueType = “numeric”,
 identifier=“http://t-sita.cesab.org/BETSI_vizInfo.jsp?trait=Body_length”),
antenna_length = as.trait(“antenna_length”,
 expectedUnit = “mm”, valueType = “numeric”,
 identifier=“http://t-sita.cesab.org/BETSI_vizInfo.jsp?trait=Antenna_length”),
metafemur_length = as.trait(“metafemur_length”,
 expectedUnit = “mm”, valueType = “numeric”,
 identifier=“http://t-sita.cesab.org/BETSI_vizInfo.jsp?trait=Femur_length”)
)
datasetStd <- standardize.traits(dataset, thesaurus = traitlist)
~~~

The function `as.thesaurus()` provides a structured object that is required by the function `standardize.traits()` (Fig. 3). Other ways of defining a ‘thesaurus’ object are documented in the package vignette and function documentation (`?as.thesaurus`). Future iterations of the R package will aim at automatising the generation of thesaurus objects from globally available ontologies. The package functions form a tool-chain where each function can be piped as an input into the next. A wrapper function `standardize( )` applies all functions sequentially, making transferring and harmonising trait data as simple as:

~~~
datasetStd <- standardize(read.csv(“path/to/data.csv”),
           thesaurus = traitlist,
           taxa = “name_correct”,
           units = “mm”
  )
~~~

Datasets that have been produced by these functions can easily be appended using the function `rbind( )` of R base, while maintaining any available metadata information as separate column entries. To merge datasets with additional information on the occurrence or measurement level, secondary data tables can be added as columns of the core datasetaccording to a unique identifier using the function `merge( )`. This enables an easy handling of data sources that originate in a relational database format.

Since the intention of the package is also to simplify the harmonisation of published trait data, the package offers direct access to trait datasets that have been released in the Public Domain or under Creative Commons licenses. We invite users and authors of datasets to add further data to the package and thereby contribute to this registry for distributed trait datasets.

## Conclusion

To serve the demand for simple ways to standardise and harmonise ecological trait data, we propose a versatile vocabulary for simple, two-dimensional datasets as well as for the exchange and handling of trait data in the context of a ‘semantic web’. With the R-package ‘traitdataform’, we also present a toolbox in R to transfer and harmonise data into this scheme.

It appears to be broad consensus that an open biodiversity science is crucial for an evidence-based decision making and conservation policy on regional and global scales. In times of increasing demand for open research data and international platforms for biodiversity data management, the development of meaningful terminologies for the standardisation of biodiversity data is more than essential: defined ontologies enable researchers to relate published datasets to each other to achieve a greater synthesis, thereby paving the way for a better mechanistic understanding of the relationship between drivers, communities and functions and providing new insights on global biodiversity patterns. Moreover it might be also a step towards a more predictive ecology as a broader set of available traits might enable more hypothesis based trait-based approaches. In terms of data science, machine-readable, ontology-based data ease the application of big-data mining and machine-learning techniques.

To date, a rich distributed body of independently published trait datasets focus on particular organism groups, ecosystem types or regions. However, these distributed data are heterogeneous in form and description and initiatives to harmonise and compile these data require significant amounts of funding and personnel. To support the long-term rewards of standardisation efforts, incentives should be sought to mitigate the cost of readying trait data for the ‘semantic web’ of biodiversity data and knowledge. This can be software tools or supporting infrastructures. The tools proposed here help to standardise trait datasets before upload to central as well as distributed data repositories. By using a constrained vocabulary with globally accessible definitions of terms, distributed trait data can be accessed more easily by other researchers and harmonised into aggregated datasets. Also, it will ease the exchange of data between databases and facilitate the development computational methods and software tools that access and handle the data, based on the standard vocabulary. We also encourage the advancement of trait thesauri into more interrelated and complete ontologies. The biggest challenge in community efforts of standardisation of traits may be the investment in consensus building which leads to an acceptance and establishment of the methodological and conceptual definitions of traits. This requires significant effort, but it returns great scientific benefit by enabling synthesis on our general understanding of biodiversity and ecosystem function.

## Acknowledgements

Thanks to all respondents to an internal online survey on trait data for the Biodiversity Exploratories project and to Matthias Biber, Kristin Bohn, Diana Bowler, Klaus Birkhofer, Runa Boeddinghaus, Catrin Westphal, Markus Fischer and Jens Kattge for comments on the manuscript drafts, the trait data standard vocabulary and the pre-releases of the R-package.

We thank the managers of the three Exploratories, Kirsten Reichel-Jung, Katrin Lorenzen, Juliane Vogt, Miriam Teuscher, and all former managers for their work in maintaining the plot and project infrastructure; Christiane Fischer and Jule Mangels for giving support through the central office, and Markus Fischer, Eduard Linsenmair, Dominik Hessenmöller, Daniel Prati, Ingo Schöning, François Buscot, Ernst-Detlef Schulze, Wolfgang W. Weisser and the late Elisabeth Kalko for their role in setting up the Biodiversity Exploratories project. The work has been partly funded by the DFG Priority Program 1374 “Infrastructure-Biodiversity-Exploratories” (DFG-Refno.).

M.J. was supported by the German Research Foundation within the framework of the Jena Experiment (FOR 1451) and by the Swiss National Science Foundation.

## Authors’ contributions

FDS, AO, CP, and NKS conceived the idea and developed the vocabulary for the trait data standard with significant contributions of MJ and GLP (forming the first tier of the author list); NKS authored the example list of traits; FDS developed the R package ‘traitdataform’; CP and FDS curated the living spreadsheet. All contributing authors appear in alphabetical order in a second tier of the author list. AG and DF implemented the vocabulary in the GFBio terminology service. All authors contributed critically to the structure and content of the manuscript and gave final approval for publication.

## Online Resources

The Appendix A contains a static excerpt of the living spreadsheet on existing trait datasets and databases, which can be found at https://goo.gl/QxzfHy.

The online reference for the Ecological Trait-data Standard Vocabulary described in this 665 paper is https://ecologicaltraitdata.github.io/ETS/, stable DOI representing all versions: 666 10.5281/zenodo.1041732.

The development website for the R-package ‘traitdataform’ is https://github.com/EcologicalTraitData/traitdataform.

Any future development of the vocabulary and the R-package is coordinated via https://github.com/EcologicalTraitData/.

